# Mammary Alveolar Proliferation and Differentiation During Early Pregnancy is Regulated by E2F5

**DOI:** 10.1101/2024.11.27.625731

**Authors:** John Vusich, Jesus Garcia-Lerena, Ryan Corfixsen, Rawan Talib, Morgan Atkins, Aishwarya V. Bhurke, Ripla Arora, Eran Andrechek

## Abstract

The E2F transcription factors are well-established cell cycle regulators, but their roles in coordinating proliferation and differentiation remain poorly understood. Here, we investigated the function of E2F5 during mammary gland development using a mammary epithelial-specific conditional knockout model. We found that E2F5 expression and chromatin binding increase during early pregnancy, coinciding with the critical window of alveolar development. Loss of E2F5 resulted in delayed alveolar expansion during early pregnancy, characterized by reduced side branching and smaller alveolar structures during early pregnancy. Mechanistically, E2F5 deletion led to reduced expression of canonical E2F target genes involved in cell cycle progression. Surprisingly, E2F5 loss also caused enrichment of luminal progenitor populations at the expense of differentiated alveolar cells, with chromatin profiling revealing substantial depletion of the repressive H3K27me3 mark at luminal progenitor-associated genes. These findings suggest that E2F5 promotes differentiation of luminal progenitors into proliferative alveolar precursors during early pregnancy. We propose that E2F5 coordinates both proliferation and differentiation by driving progenitor cells to differentiate into mature luminal cells. The dual function of E2F5 in mammary development distinguishes it from classical cell cycle regulators and positions it as a critical coordinator of the developmental transitions required for lactation.

## Introduction

The mammary gland sets mammals apart from other vertebrates through its specialized function of synthesizing and secreting milk to nourish developing offspring^1^. Mammary glands exhibit remarkable regenerative capacity, undergoing substantial structural and functional transformations across a female’s lifespan, particularly during puberty, pregnancy, and lactation^2^. Following puberty, the adult mammary gland features a bilayered architecture, with luminal epithelial cells positioned apically to enclose the central lumen and basal/myoepithelial cells forming an exterior layer adjacent to the basement membrane^3^. Within the luminal compartment, there exists considerable cellular heterogeneity, including actively dividing luminal progenitor (LP) cells and hormone-receptor (HR) positive (HR^+^) mature luminal (ML) cells^4^. The LP population itself comprises a HR^−^ subset alongside a minor HR^+^ fraction, with the latter thought to give rise to the HR^+^ ML lineage^4,5^. At the onset of pregnancy, progesterone and prolactin signaling trigger extensive tissue remodeling requiring coordinated interactions among basal cells, hormone-sensing HR+ luminal progenitor cells, and HR- alveolar luminal cells^6^. This alveologenesis developmental program proceeds through two sequential phases: first, a proliferative phase characterized by rapid epithelial expansion that leads to ductal expansion and generates alveolar structures, followed by a differentiation phase during which alveolar progenitors mature into functional, milk-secreting alveolar cells^6,7^. Throughout these dynamic morphogenic transitions, precise transcriptional control of proliferation and differentiation is paramount to ensure proper mammary function^8^.

Luminal progenitor cells are epigenetically primed to undergo proliferation and differentiation in response to pregnancy hormones, particularly progesterone and prolactin^9,10^. This epigenetic priming extends to the parous mammary gland, enabling more efficient responses to subsequent pregnancies^11^. Notch^12,13^ signaling and the transcription factors GATA3^14,15^, FOXA1^16^, and FOXP1^17^ are indispensable for postnatal mammary development and establishment of the luminal lineage during ductal morphogenesis. During pregnancy, progesterone^18–20^, RANK^21,22^, and PRLR/JAK/STAT^23–25^ signaling are critical for alveologenesis. STAT6 promotes cellular proliferation and survival in early pregnancy^26^, while STAT5^25,27^, ELF5^13,22,28^, and EHF^29^ drive establishment of the secretory alveolar lineage. Despite these advances, the molecular mechanisms that regulate cellular proliferation and differentiation during the transition from the adult virgin state to pregnancy in the mammary gland remain incompletely understood.

The E2F family of transcription factors, through regulation by cyclins and the RB family of proteins, play a pivotal role in controlling cell cycle progression^30,31^. E2F transcription factors are traditionally comprised of transcriptional activators (E2F1-3) and repressors (E2F4-8), with each subfamily playing distinct roles in regulating gene expression during various cellular processes. Beyond their canonical role in cell cycle control, E2Fs have been found to regulate additional processes, including cellular differentiation^32,33^, apoptosis^34,35^, DNA damage response^36,37^, and metabolism^38,39^. Emerging evidence indicates that E2Fs can function independently of RB regulation during development, suggesting additional layers of transcriptional control beyond canonical cell cycle pathways^40^. In the context of mammary gland development, increased expression and pathway activity of E2Fs during proliferative phases, coupled with delayed development observed in mice with global knockouts of various E2F family members, suggests an important role for E2Fs in regulating mammary gland development^41^.

The specific role of E2F5 in mammary gland development has been understudied, largely due to the embryonic lethality associated with E2F4/5 double knockout and postnatal lethality of global E2F5 knockout^42,43^. Global E2F5 knockout in mice leads to postnatal lethality due to hydrocephalus resulting from a differentiation, rather than proliferation, defect in the choroid plexus^42^. While E2F5 is traditionally classified as a transcriptional repressor, emerging evidence demonstrates that E2F5 can function as a transcriptional activator in specific developmental contexts. For example, during multiciliate cell development, E2F5 forms a ternary complex with Multicilin and Dp1 to activate genes required for centriole biogenesis^44^. E2F5 has been implicated in regulating developmental differentiation programs across several organ systems, including the central nervous^42,45^, digestive^46^, male reproductive^47^, and respiratory systems^44^. Given E2F5’s role in developmental differentiation and our recent findings that E2F5 plays a role in mammary development during puberty and is a putative tumor suppressor gene in the mammary gland^48^, we sought to investigate whether E2F5 regulates mammary epithelial differentiation during pregnancy.

To investigate the role of E2F5 in mammary gland development during pregnancy, we developed a mouse model that allows mammary conditional deletion of the *E2f5* gene. We analyzed expression and genomic occupancy of E2F5 in primary mouse mammary epithelial cells (MECs) during pregnancy compared to the adult virgin state. Both E2F5 expression and chromatin binding increased from virgin to pregnancy, suggesting an important role in regulating the developing epithelium during pregnancy. Utilizing our E2F5 conditional knockout (CKO) mice, we found that loss of E2F5 resulted in delayed alveolar development during early pregnancy, accompanied by reduced proliferation, decreased expression of mature luminal markers, and enrichment of luminal progenitor and mammary stemness pathways. These findings suggest that E2F5 promotes proliferative expansion and terminal differentiation of mammary epithelial cells during pregnancy.

## Results

### Expression and activity of E2F5 increases in the mammary gland during pregnancy

We have previously examined the role of various E2Fs in various cycles of mammary development, demonstrating a role for E2F1,3 and 4 in puberty, E2F4 in lactation, and E2F3 during involution. Given that we recently observed pubertal ductal elongation delays with a mammary specific deletion of E2F5, we have analyzed E2F5 expression using publicly available RNA-seq data from mammary glands across multiple developmental stages. This included virgin, pregnancy day 6, pregnancy day 18, lactation day 1, and lactation day 10. E2F5 expression in whole mammary glands remained relatively consistent across these conditions (Figure 1A). To further examine cell-type-specific expression, we analyzed publicly available RNA-seq data from isolated basal and luminal cells at virgin, pregnancy day 18.5, and lactation day 2 stages. E2F5 expression increased during pregnancy, particularly in luminal cells, relative to the virgin and lactating stages (Figure 1B-C).

**Figure 1.**
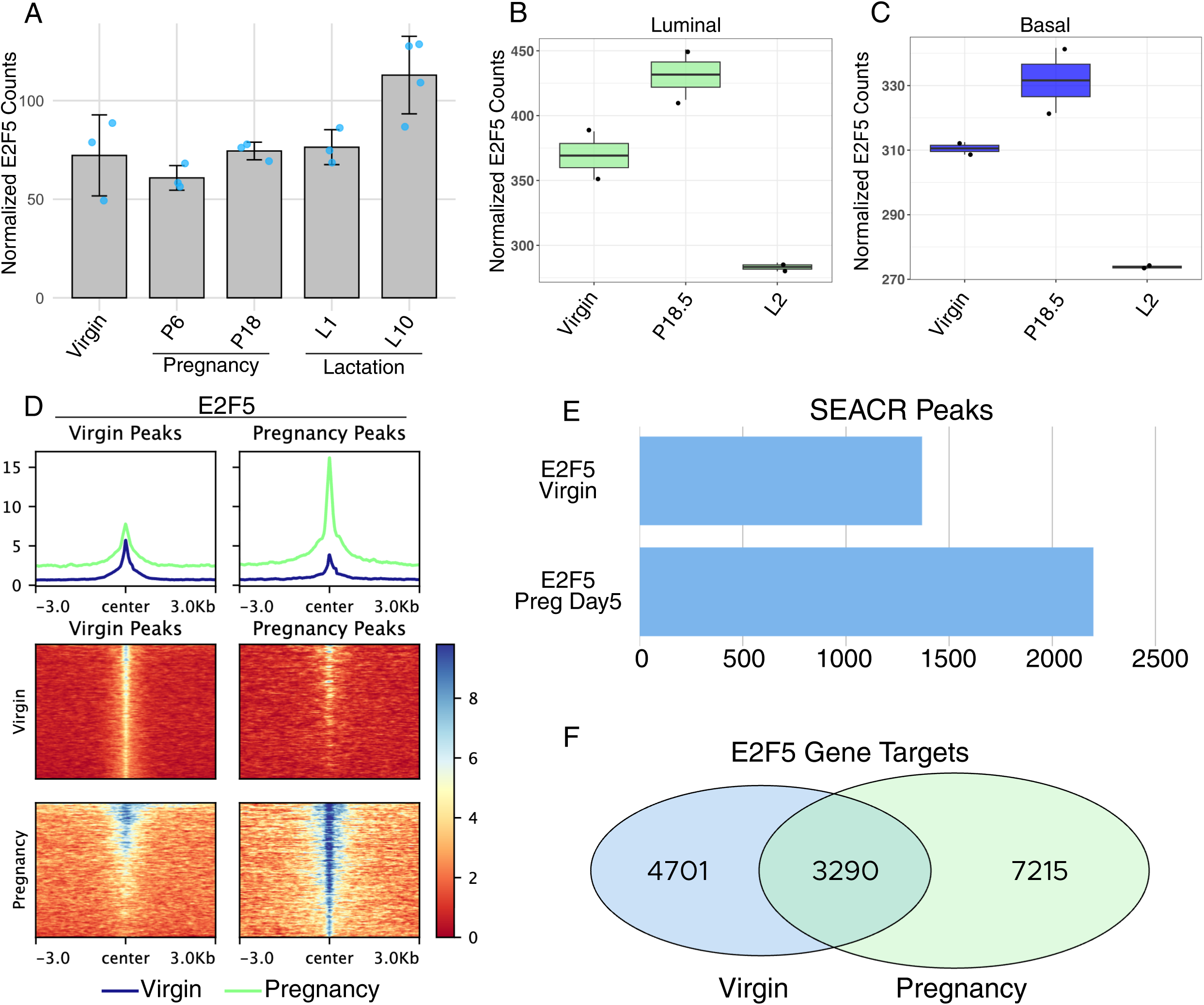
E2F5 expression and chromatin binding increase during pregnancy in the mammary gland. (A) Normalized E2F5 expression in whole mammary glands across developmental stages: virgin, pregnancy day 6 (P6), pregnancy day 18 (P18), lactation day 1 (L1), and lactation day 10 (L10). (B, C) Normalized E2F5 expression in sorted luminal (B) and basal (C) mammary epithelial cells at virgin, pregnancy day 18.5 (P18.5), and lactation day 2 (L2) stages. (D) E2F5 CUT&RUN signal in mammary epithelial cells from virgin (left) and pregnancy day 5 (right) mice. Top: average profile plots aligned to peak centers; bottom: corresponding heatmaps for peaks called in virgin (1,369 peaks) and pregnancy day 5 (2,199 peaks) samples. (E) Number of E2F5 peaks identified by SEACR in virgin and pregnancy day 5 mammary epithelial cells. (F) Venn diagram showing overlap and unique E2F5 target genes in virgin and pregnancy day 5 samples.

Next, we performed CUT&RUN to identify E2F5 target genes in primary mammary epithelial cells from virgin mice and mice on day 5 of pregnancy. CUT&RUN analysis revealed increased E2F5 chromatin binding during early pregnancy. E2F5 binding intensity was higher in pregnancy day 5 mammary epithelial cells compared to virgin cells at both virgin and pregnancy-specific peak regions (Figure 1D). Consistent with this observation, SEACR peak calling identified substantially more E2F5 binding sites in pregnancy day 5 cells (2,199 peaks) compared to virgin cells (1,369 peaks) (Figure 1E). Analysis of E2F5 target genes using ChIPseeker revealed 3,290 shared target genes between virgin and pregnancy day 5 conditions, while 4,701 and 7,215 genes were uniquely targeted in virgin and pregnancy day 5 cells, respectively (Figure 1F). These data indicate that E2F5 engages a broader set of genomic targets during early pregnancy.

### E2F5 chromatin binding increases during early pregnancy in concordance with H3K4me3 occupancy

To investigate the relationship between E2F5 binding and transcriptional activity, we performed CUT&RUN for H3K4me3, a histone mark associated with active gene promoters and enhancers. Genome-wide analysis revealed high concordance between E2F5 and H3K4me3 occupancy in both virgin and pregnancy day 5 mammary epithelial cells, with both signals enriched near transcription start sites (Figure 2A). Notably, while the target genes were largely consistent between conditions, both E2F5 and H3K4me3 binding intensity increased substantially during early pregnancy (Figure 2A-B). An UpSet plot analysis confirmed extensive overlap between E2F5- and H3K4me3-marked genes, with the majority of genes co-occupied by both marks in each condition (Figure 2B). This coordinated increase in binding intensity was evident at individual pregnancy-associated mammary development genes, where E2F5 and H3K4me3 signals were concordantly elevated at pregnancy day 5 compared to virgin cells (Figure 2C). These findings suggest that E2F5 activity during early pregnancy is associated with increased transcriptional potential at a consistent set of target genes rather than engagement of entirely new genomic loci.

**Figure 2.**
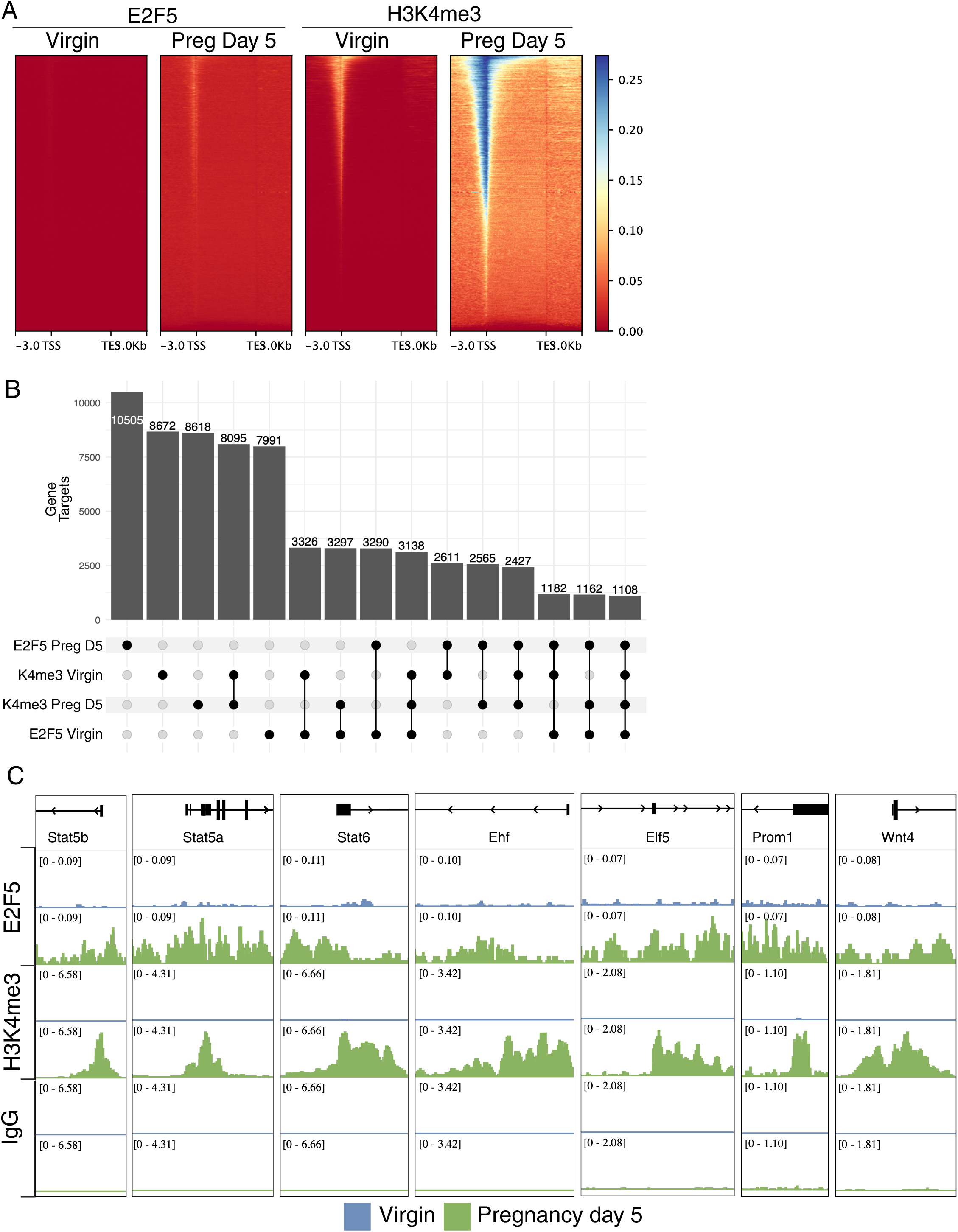
E2F5 chromatin binding increases during early pregnancy in concordance with H3K4me3 occupancy. (A) deepTools heatmaps showing E2F5 and H3K4me3 CUT&RUN signal across all annotated mm39 genes (±3 kb from the transcription start site [TSS] and transcription end site [TES]) in mammary epithelial cells from virgin and pregnancy day 5 mice. Increased E2F5 binding during pregnancy coincides with an increased H3K4me3 signal near the TSS. (B) UpSet plot showing intersections of gene targets bound by E2F5 and/or marked by H3K4me3 across virgin and pregnancy day 5 samples. Bar heights represent the number of genes in each intersection. (C) IGV browser snapshots showing E2F5 and H3K4me3 signals at representative pregnancy-associated genes, illustrating concordant occupancy.

### Loss of E2f5 in the mammary gland impairs alveolar development during early pregnancy

To investigate the functional role of E2F5 in mammary gland development, we generated mammary epithelial-specific conditional knockout mice by crossing MMTV-Cre with E2F5 floxed mice (Figure 3A). We compared mammary gland morphology in MMTV-Cre E2F5^F/F^ mice and E2F5^F/F^ littermate controls at virgin and pregnancy day 5 stages using wholemount and histological analyses. Virgin mammary glands from 12-week-old mice showed normal ductal architecture in both control and E2F5 conditional knockout animals, indicating that E2F5 is dispensable for epithelial filling of the mammary fat pad (Figure 3B-E). In contrast, mammary glands from E2F5 conditional knockout mice at pregnancy day 5 displayed impaired alveolar development compared to controls, with a slight delay of expansion (Figure 3F-I). Quantification of wholemounts revealed that E2F5 conditional knockout mice had significantly reduced pixel density and area of lateral side branches (Figure 3J-K), while the hand-counted number of alveoli on lateral side branches was not significantly different between genotypes (Figure 3L). These data demonstrate that E2F5 is required for proper alveolar development during early pregnancy but is dispensable for virgin mammary gland morphogenesis.

**Figure 3.**
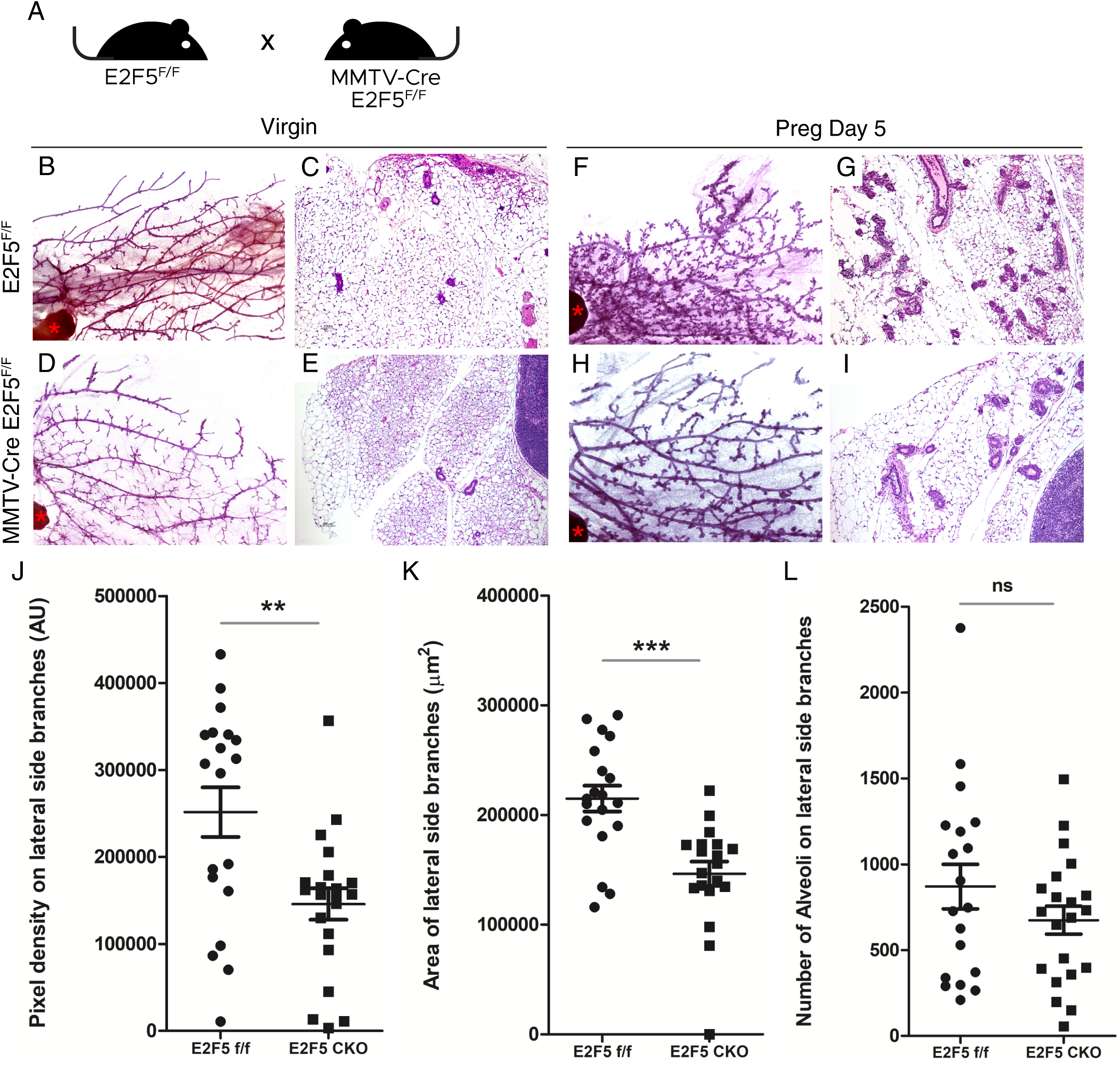
Loss of E2F5 in the mammary epithelium delays alveolar development during early pregnancy. (A) Breeding strategy to generate E2F5 mammary conditional knockout mice (MMTV-Cre E2F5^F/F^) and E2F5^F/F^ control littermates. (B–I) Representative whole mounts (B, D, F, H) and H&E-stained FFPE sections (C, E, G, I) of mammary glands from 12-week-old virgin females (B–E) and day 5 pregnant females (F–I). E2F5^F/F^ mice are shown in the top row (B, C, F, G) and MMTV-Cre E2F5^F/F^ mice in the bottom row (D, E, H, I). Whole mounts highlight ductal and alveolar morphology, while histological sections show epithelial structure within the fat pad. (J–L) Quantification from whole mounts: (J) pixel density of lateral side branches (p = 0.0025), (K) area of lateral side branches (p < 0.001), and (L) number of alveoli on lateral side branches (ns, not significant). Each data point represents one animal; error bars show mean ± SEM. Significance determined by unpaired two-tailed t-test.

To further characterize the mammary gland phenotype in E2F5 conditional knockout mice, we performed confocal imaging of intact Hoechst-stained wholemounts from pregnancy day 5 animals. Compared to control littermates, about 50% of the MMTV-Cre E2F5^F/F^ mammary glands appeared smaller overall with altered epithelial architecture (Figure 4A-D). While the conditional knockout glands displayed an irregular branching pattern with apparent increased branching density, the developing alveolar structures (arrows) appeared smaller, and the ductal diameter (arrowheads) appeared larger relative to controls (Figure 4E-H). Additionally, the total amount of epithelium within the mammary fat pad appeared to be reduced in E2F5 conditional knockout glands. Together with the wholemount analysis, these findings indicate that E2F5 regulates multiple aspects of mammary epithelial development during early pregnancy, including alveolar size, ductal morphology, and overall epithelial content.

**Figure 4.**
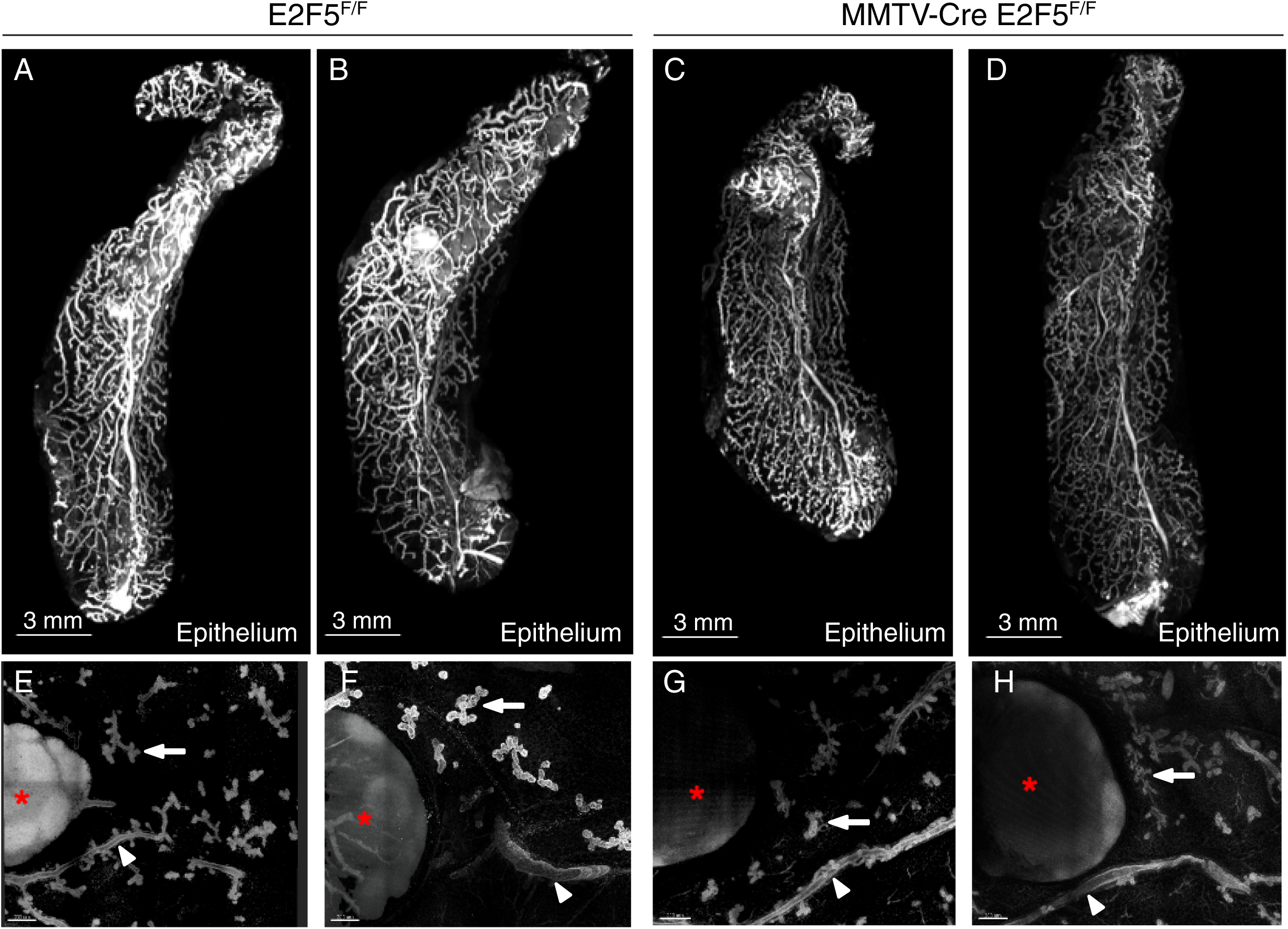
Loss of E2F5 leads to smaller mammary glands with an irregular branching pattern. (A-D) Confocal imaging of E2F5^F/F^ (A-B) and MMTV-Cre E2F5^F/F^ (C-D) whole mammary glands, showing epithelium (Hoechst-stained cells) in white. (E-H) High-resolution cross sections from confocal images of E2F5^F/F^ (E-F) and MMTV-Cre E2F5^F/F^ (G-H) whole mammary glands, showing epithelium (Hoechst-stained cells) in white.

### Loss of E2F5 reduces proliferation-associated gene expression in mammary epithelial cells during early pregnancy

To investigate the transcriptional consequences of E2F5 loss that are involved in the delayed pregnancy development, we isolated primary mammary epithelial cells from pregnancy day 5 E2F5^F/F^ control and MMTV-Cre E2F5^F/F^ mice using magnetic-activated cell sorting and performed bulk RNA-seq (Figure 5A). Gene set enrichment analysis revealed significant downregulation of proliferation-associated pathways in E2F5 conditional knockout cells, including Hallmark E2F targets, Hallmark G2M checkpoint, Kong E2F3 targets, and genes upregulated upon acute RB1 loss of function (Figure 5B-C). In contrast, we observed upregulation of immune and tumor microenvironment-associated pathways, including epithelial-mesenchymal transition, TNFα signaling via NFκB, interferon gamma response, and complement pathway genes (Figure 5B). To confirm that these downregulated genes are direct E2F5 targets, we examined E2F5 and H3K4me3 occupancy at representative G2M checkpoint genes (Ccna2, Ccnb1, Cdk1) and E2f1 using our pregnancy day 5 CUT&RUN data. All four genes showed increased E2F5 and H3K4me3 binding during early pregnancy compared to virgin cells, consistent with their regulation by E2F5 (Figure 5D). To assess proliferation at the tissue level, we performed immunohistochemical staining for PCNA, a marker of proliferating cells. While PCNA staining intensity appeared similar between genotypes, E2F5 conditional knockout mammary glands exhibited fewer PCNA-positive clusters, suggesting reduced proliferative capacity in developing alveolar structures (Figure 5E-F). Together, these findings demonstrate that E2F5 promotes expression of proliferation-associated genes and is required for normal proliferative expansion during early pregnancy mammary gland development.

**Figure 5.**
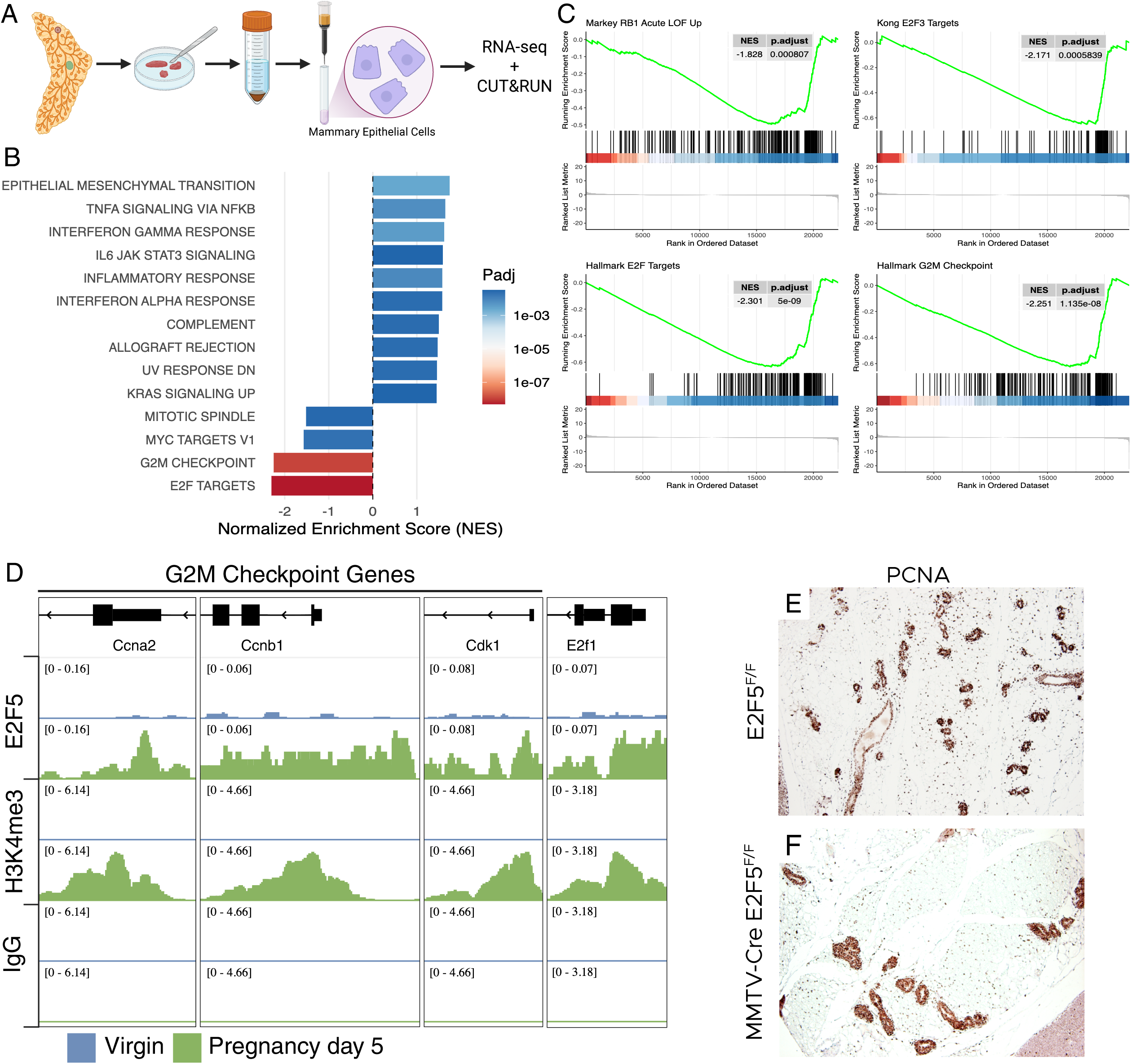
Loss of E2F5 reduces E2F target gene expression and proliferation in mammary epithelial cells. (A) Schematic of experimental workflow. Mammary glands from pregnancy day 5 E2F5^F/F^ control and MMTV-Cre E2F5^F/F^ conditional knockout mice were dissected, dissociated into single-cell suspensions, and subjected to magnetic-activated cell sorting (MACS) depletion using antibodies against CD45, CD31, Ter119, and CD34 to remove non-epithelial cells. Flow-through fractions enriched for mammary epithelial cells (MECs) were collected for bulk RNA-seq and CUT&RUN. (B) Gene set enrichment analysis (GSEA) bar plot showing significantly altered hallmark gene sets. (C) Individual GSEA plots showing significant downregulation of the Hallmark E2F targets, Kong E2F3 targets, Hallmark G2M checkpoint, and RB1 acute loss-of-function upregulated gene sets in E2F5 CKO MECs compared to E2F5 F/F controls. (D) IGV plots showing occupancy of E2F5, H3K4me3, and IgG in Virgin (blue) and Pregnancy Day 5 (green) MECs at the promoter regions of G2M-associated genes (Ccna2, Ccnb1, and Cdk1) and E2f1. (E, F) Immunohistochemical staining for PCNA, a proliferation marker, in FFPE mammary gland sections from pregnancy day 5 E2F5^F/F^ (E) and MMTV-Cre E2F5^F/F^ (F) mice.

### E2F5 is largely dispensable for late-pregnancy and subsequent-pregnancy mammary gland development

We examined mammary gland development at later stages to determine the temporal requirement for E2F5. At pregnancy day 14, both control and E2F5 conditional knockout mice displayed extensive alveolar development with minimal morphological differences (Figure 6A-D). Prior reports that discussed epigenetic priming^10,11,49^ led us to examine the second pregnancy at day 5. This revealed that mammary glands from both genotypes showed comparable architecture and robust alveolar development with less pronounced differences than the first pregnancy (Figure 6E-H). Gene set enrichment analysis revealed up-regulation of genes associated with post-pregnancy chromatin remodeling in E2F5 conditional knockout cells (Figure 6I), suggesting potential compensatory mechanisms during later developmental stages.

**Figure 6.**
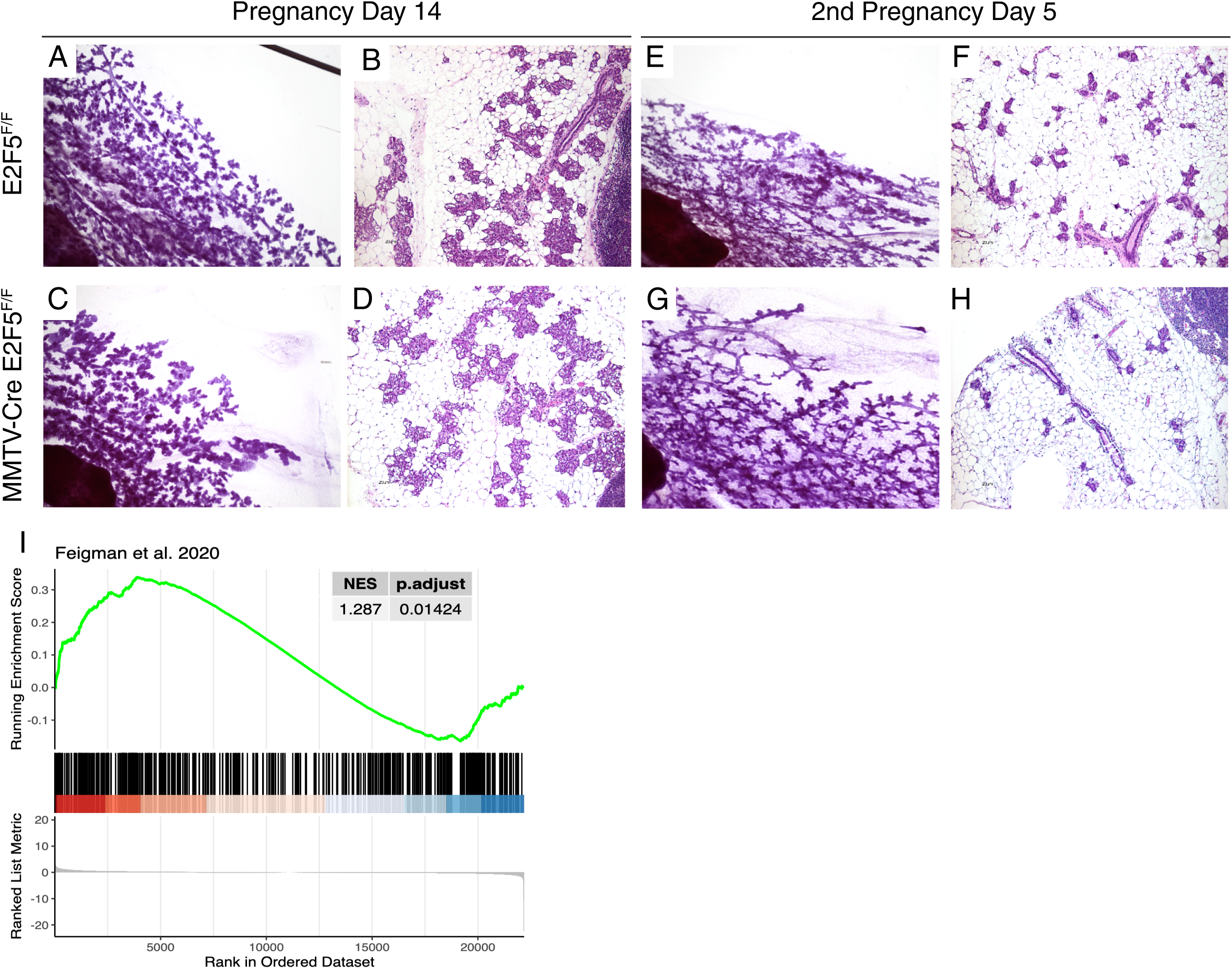
Loss of E2F5 alters ductal branching and alveolar morphology during mid-pregnancy and second pregnancy. (A–H) Representative whole mounts (A, C, E, G) and H&E-stained FFPE sections (B, D, F, H) of mammary glands from E2F5^F/F^ control (top row) and MMTV-Cre E2F5^F/F^ conditional knockout (bottom row) mice. Samples were collected on pregnancy day 14 (A–D) and second pregnancy day 5 (E–H). Whole mounts highlight ductal branching and alveolar development, while histological sections show epithelial organization within the fat pad. (I) GSEA using differentially expressed genes that gain H3K27ac post-pregnancy from Feigman et al. 2020^49^, showing upregulation of the gene set in MMTV-Cre E2F5^F/F^ versus E2F5^F/F^ control.

### Loss of E2F5 impairs luminal progenitor differentiation and enriches progenitor cell populations

To investigate the cellular basis of impaired alveolar development in E2F5 conditional knockout mice, we performed unsupervised hierarchical clustering of the top 30 most variable genes from our pregnancy day 5 RNA-seq data. This analysis revealed distinct gene expression patterns between genotypes, with E2F5 conditional knockout samples showing elevated expression of luminal progenitor-associated genes, including Kit, a marker of luminal progenitor cells (Figure 7A). Gene set enrichment analysis confirmed this shift in cellular identity, revealing significant upregulation of mammary luminal progenitor and mammary stem cell gene signatures in E2F5 conditional knockout cells, coupled with downregulation of mammary luminal mature gene signatures (Figure 7B). To investigate the chromatin landscape underlying these changes, we performed CUT&RUN for three key histone marks, including H3K4me3 and H3K27ac (activating marks) and H3K27me3 (repressive mark) in E2F5 conditional knockout and control mammary epithelial cells. Examining luminal progenitor driver genes, including Kit, Hey1, Foxi1, and Cd14, we observed that activating histone marks were maintained at comparable levels between genotypes, while the repressive H3K27me3 mark was substantially depleted in E2F5 conditional knockout cells compared to controls (Figure 7C). These chromatin changes suggest that loss of E2F5 leads to transcriptional de-repression of luminal progenitor genes, favoring progenitor cell maintenance over differentiation. Together, these findings indicate that E2F5 promotes luminal progenitor differentiation into alveolar cells during early pregnancy, and its loss results in enrichment of progenitor populations at the expense of differentiated alveolar cells (Figure 7D).

**Figure 7.**
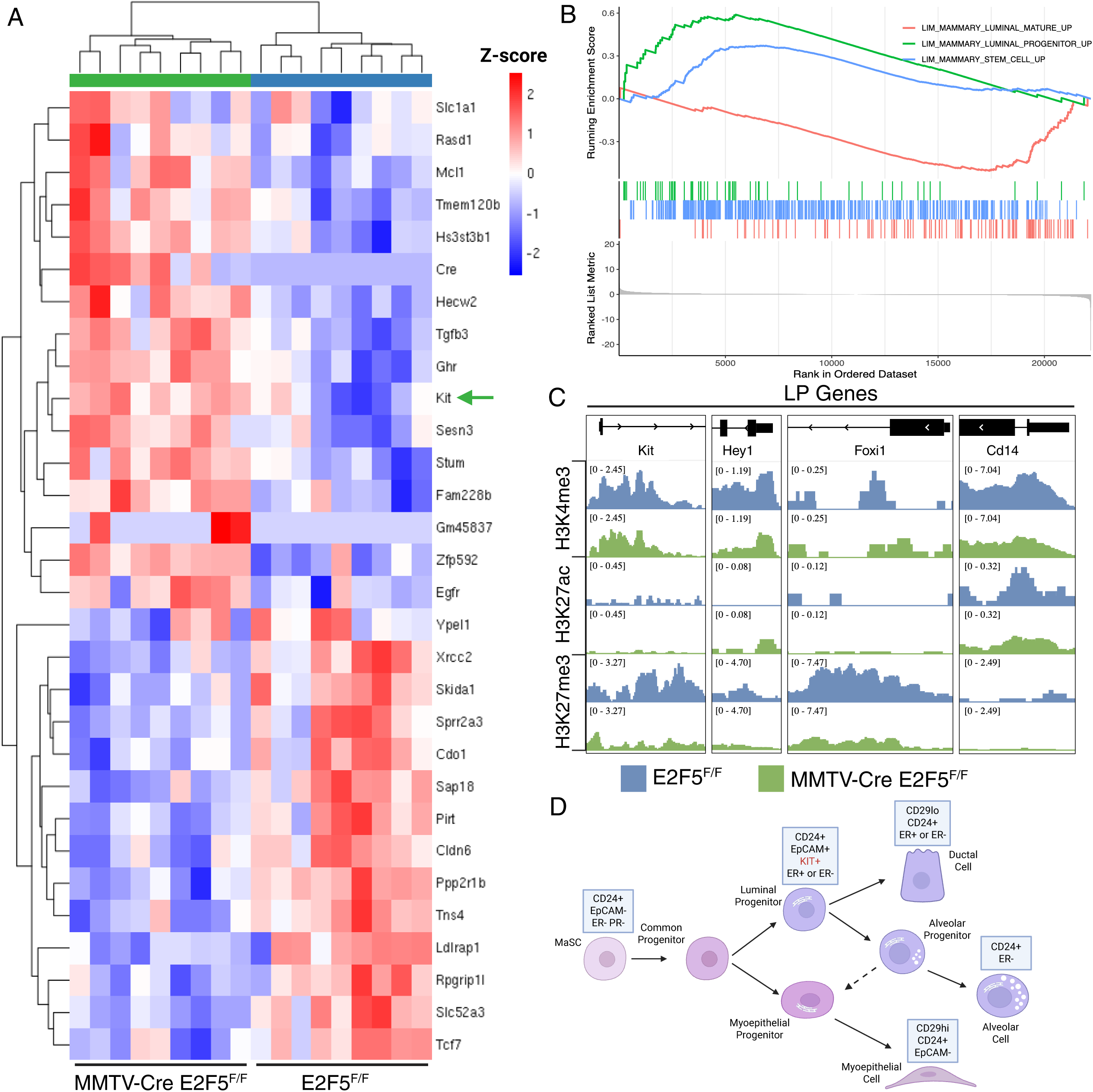
Loss of E2F5 enriches luminal progenitor and mammary stem cell populations at the expense of differentiated alveolar cells. (A) Heatmap of differentially expressed genes from pregnancy day 5 mammary epithelial cells showing altered expression of lineage-associated genes in MMTV-Cre E2F5^F/F^ conditional knockout (CKO) mice compared to E2F5^F/F^ controls. Genes associated with luminal progenitor identity (e.g., Kit) are enriched in CKO MECs. (B) Combined gene set enrichment analysis (GSEA) plot showing downregulation of the mammary luminal mature signature (red) and upregulation of mammary luminal progenitor (blue) and mammary stem cell (green) signatures in E2F5 CKO compared to controls. (C) Proposed model illustrating that loss of E2F5 leads to enrichment of cKit+ luminal progenitor cells, impairing their differentiation into alveolar cells and reducing alveolar development during pregnancy.

## Discussion

In this study, we investigated the role of E2F5 in mammary gland development during pregnancy. We found that E2F5 expression and chromatin binding activity increase during early pregnancy, coinciding with the critical window of alveolar development. Using a mammary epithelial-specific conditional knockout model, we demonstrated that E2F5 is required for proper alveolar expansion during early pregnancy but is dispensable for both virgin gland development and later pregnancy stages. Loss of E2F5 resulted in reduced expression of proliferation-associated genes and impaired alveolar development. Mechanistically, E2F5 deletion led to enrichment of luminal progenitor populations through chromatin-mediated up-regulation of mammary luminal progenitor-associated genes, suggesting that E2F5 promotes luminal progenitor differentiation into alveolar cells during early pregnancy.

The temporal specificity of the E2F5 requirement is particularly striking. E2F5 conditional knockout mice displayed delayed ductal development during puberty, though this phenotype eventually resolved as glands matured to the adult virgin state. Similarly, during pregnancy, the E2F5 conditional knockout mice exhibited delayed alveolar development at pregnancy day 4, with reduced side branching and smaller alveolar structures. This phenotype largely resolved by pregnancy day 14, indicating that compensatory mechanisms can rescue alveolar development at both stages. This stage-specific during these two critical developmental windows, puberty and early pregnancy, aligns with the dynamic expression pattern we observed, where E2F5 levels increase during pregnancy, particularly in luminal cells. The coordination between E2F5 expression, chromatin binding, and the H3K4me3 active promoter mark suggests that E2F5 functions as part of a transcriptional program that drives the proliferative expansion necessary for ductal elongation during pregnancy and alveolar development during pregnancy. This is consistent with prior work demonstrating recruitment of Stat5 followed by H3K4me3 marks and transcriptional activation in the mammary gland^50^.

Our finding that E2F5 loss reduces expression of canonical E2F target genes, including cell cycle regulators such as Ccna2, Ccnb1, Cdk1, and E2f1, is consistent with E2F5’s known role in cell cycle regulation^31^. E2F5 has historically been characterized as a repressive transcription factor, often antagonizing the activating E2Fs (E2F1-3)^51^. Our data support the notion that E2Fs have non-cell cycle related functions, here in the context of mammary gland development. The increased E2F5 chromatin binding at proliferation-associated genes during pregnancy, coupled with the reduction in these genes upon E2F5 loss, suggests that E2F5 may function as a transcriptional activator, promoting expansion of the luminal epithelial compartment in the mammary gland during pregnancy. Indeed, this is consistent with the upregulation of many genes with the overexpression of E2F5 when used to generate a genomic signature for E2F5 activity^48^. Alternatively, E2F5 may be required to maintain chromatin accessibility at these loci, facilitating activation by other E2F family members or transcription factors. Indeed, E2Fs have been associated with chromatin accessibility in prior work. For instance, E2Fs in conjunction with Stat3 have been demonstrated to maintain chromatin accessibility in a glioblastoma model^52^. Further, in a developmental context in *Drosophila*, disruption of E2F binding sites in PGK was associated with a marked reduction in chromatin accessibility^53^. However, recent work has shown that E2Fs can activate proliferation associated genes without changing chromatin accessibility^54^. The concurrent increase in both E2F5 and H3K4me3 at target genes during pregnancy supports either model.

The most intriguing finding from our study is that E2F5 loss leads to enrichment of luminal progenitor populations at the expense of differentiated alveolar cells, which is manifested in the observed phenotype. Gene expression analysis revealed up-regulation of luminal progenitor and mammary stem cell signatures, including increased expression of *Kit*, a well-established marker of luminal progenitors^5,55^. Chromatin profiling revealed that luminal progenitor-associated genes showed substantial depletion of the repressive H3K27me3 mark in E2F5 conditional knockout cells, while activating marks (H3K4me3 and H3K27ac) remained largely unchanged. This chromatin landscape is permissive for gene expression, potentially explaining the transcriptional upregulation of progenitor genes. These findings suggest that E2F5 normally promotes the differentiation of luminal progenitors into proliferating alveolar precursors, and its absence results in a differentiation block that maintains cells in a progenitor-like state. In support of this, it has been demonstrated that E2Fs regulate EZH2 expression and that this is key in maintaining H3K27 methylation^56^. Indeed, a study in glioblastoma demonstrated that E2F7 promoted EZH2 expression, leading to an increase in H3K27me3 levels^57^.

Here we have shown that E2F5 has regulated both proliferation and differentiation simultaneously. This raises the question of how E2F5 might coordinate the regulation of these divergent transcriptional programs. One possibility is that E2F5 promotes the expression of genes required for proliferative expansion while simultaneously repressing progenitor maintenance. The apparent contradiction of reduced proliferation yet enriched progenitor populations in E2F5 conditional knockout glands can be reconciled by considering the heterogeneity within the luminal progenitor compartment. Recent work has identified hormone-receptor (HR) negative luminal progenitors (HR-LPs) and hormone-sensing luminal progenitors (HR+LPs) in the mammary gland, both of which are proliferative^58,59^. Notably, while HR+LPs exhibit the highest proliferation rates, they do not expand numerically, instead likely undergoing rapid asymmetric cell division and differentiation into mature luminal cells^59^. In contrast, HR-LPs maintain and expand their population during proliferation, remaining poised for alveolar differentiation during pregnancy. We propose that E2F5 promotes proliferation and differentiation in response to pregnancy hormones, primarily progesterone, in the HR+LP cells. E2F5 may regulate distinct gene programs in different cellular contexts, functioning as an activator at proliferation genes and a repressor at progenitor genes, potentially through interactions with different cofactors or chromatin modifiers. Given that E2F5 is known to associate with cofactors including Smad4 to regulate genes such as c-Myc^60^, and that Myc is known to regulate mammary development and function^61^, the regulation of proliferation and differentiation by E2F5 in pregnancy illustrates a unique transcriptional regulatory role amongst the E2F transcription factors.

The observation that E2F5 conditional knockout mammary glands had resolved compared to the control counterparts by later stages of pregnancy raises important questions about redundancy within the E2F family. Other E2F family members, particularly E2F1-4, are expressed in the mammary gland and may compensate for E2F5 loss as pregnancy progresses. Indeed, prior work demonstrated that while loss of E2F2 alone had no effect on mammary gland development, double knockouts of E2F1 and E2F2 or E2F3 and E2F2 had much more severe ductal elongation delays^41,62^. Additionally, pregnancy hormones, particularly prolactin, increase dramatically as pregnancy advances and may activate alternative pathways that bypass the E2F5 requirement. The upregulation of genes associated with post-pregnancy chromatin remodeling in second pregnancy E2F5 conditional knockout mice suggests that epigenetic reprogramming after the first pregnancy may also contribute to compensation. Indeed, this is reminiscent of the epigenetic memory associated with pregnancy^11,49^. Understanding these compensatory mechanisms will be important in future work for determining whether E2F5 has unique functions or is part of a more redundant system.

## Conclusion

In conclusion, we have identified E2F5 as a stage-specific regulator of mammary gland development that promotes luminal progenitor differentiation and proliferative expansion during early pregnancy. Our findings reveal an unexpected function for E2F5 beyond its canonical role in cell cycle regulation and highlight the importance of coordinating proliferation and differentiation during developmental transitions. Further investigation of the mechanisms by which E2F5 regulates chromatin state and cell fate decisions will deepen our understanding of mammary gland biology and may inform strategies for addressing breast cancer.

## Methods

### Mammary Gland Isolation

Mammary glands were harvested and dissociated as previously described. In brief, inguinal mammary glands from each mouse were dissected, transferred to a sterile 50 mL conical tube, and minced with scissors. Minced glands were incubated in pre-warmed Dissociation Solution I (DS1; DMEM/F12 HEPES supplemented with 1 mg/mL Collagenase D, 1 mg/mL Hyaluronidase, and 1.875% BSA) and shaken at 37 °C for 2 hours to enable enzymatic digestion. Tubes were vortexed briefly every 15 minutes to facilitate dissociation. Following digestion, samples were vortexed for 5 seconds and transferred to clean 15 mL conical tubes. Cell suspensions were centrifuged at 350 × g for 5 minutes, and pellets were resuspended in 2 mL of pre-warmed TrypLE Express. Samples were shaken at 37 °C for 2 minutes, and the reaction was quenched with 10 mL of ice-cold HF (HBSS containing 5% FBS). Pelleted cells were collected at 350 × g for 5 minutes, resuspended in 2 mL of pre-warmed Dissociation Solution II (DS2; DMEM/F12 HEPES supplemented with 5 mg/mL Dispase II and ∼330 U/mL recombinant DNase I), and incubated with shaking at 37 °C for 2 minutes. Digestions were quenched with 10 mL of cold HF, and resulting suspensions were filtered through a 40 µm cell strainer. When necessary, red blood cells were lysed using a 1:4 mixture of cold HF:NH Cl prior to downstream processing.

### Histology

Mammary glands were fixed in 10% formalin, embedded in paraffin, and sectioned. Tissue sections were stained with haematoxylin and eosin (H&E). For wholemounts, mammary glands were placed on a slide and fixed in Carnoy’s solution overnight at room temperature before staining in carmine alum.

### Lineage Depletion and Epithelial Cell Enrichment

Following dissociation, single-cell suspensions were resuspended in MACS buffer (DPBS containing 0.5% BSA and 2 mM EDTA) and incubated with TruStain FcX™ PLUS anti-mouse CD16/32 (BioLegend, #156603) for 10 minutes on ice at 0.25 µg per 10□ cells to block Fc receptors. Cells were pelleted (350 × g, 5 minutes), resuspended in fresh MACS buffer, and incubated with a biotinylated lineage antibody cocktail. After incubating on ice for 10–15 minutes, the cells were washed twice with MACS buffer and then resuspended in 70 µL of MACS buffer per sample. Anti-biotin microbeads (Miltenyi Biotec, #130-090-485) were added at 20 µL per sample, mixed thoroughly, and incubated for 15 minutes on ice. Labeled cells were washed with 2 mL MACS buffer and resuspended in 500 µL for magnetic separation. Samples were applied to LS separation columns (Miltenyi Biotec, #130-042-401) mounted on a magnetic rack. Columns were washed three times with 3 mL MACS buffer, and the flow-through, representing the lineage-negative mammary epithelial cell population, was collected. These epithelial-enriched fractions were used for downstream RNA-seq and CUT&RUN experiments.

### Antibodies

Antibodies for lineage depletion: anti-CD45 (eBioscience, #13-0451-85; RRID:AB_466447), anti-CD31 (eBioscience, #13-0311-85; RRID:AB_466421), anti-Ter119 (eBioscience, #13-5921-85; RRID:AB_466798), and anti-CD34 (eBioscience, #13-0341-82; RRID:AB_466425) were used at a 1:100 dilution. Antibodies for immunohistochemistry: rabbit monoclonal anti-PCNA (Clone D3H8P; Cell Signaling #13110-15; RRID:AB_2636979) was used at a 1:4000 dilution. Antibodies for CUT&RUN: rabbit IgG (Epicypher #13-0042; RRID:AB_2923178), rabbit monoclonal anti-H3K27me3 (Invitrogen #MA5-11198; RRID:AB_1100074), rabbit monoclonal anti-H3K4me3 (Epicypher #13-0041; RRID:AB_3076423), rabbit monoclonal anti-H3K27ac (Epicypher #130059; RRID:AB_3712075), and rabbit polyclonal anti-E2F5 (Invitrogen #PA585578; RRID:AB_2792718) were used at 0.5 ug per sample replicate in 50 uL of volume.

### CUT&RUN

CUT&RUN was performed using the CUTANA ChIC/CUT&RUN kit (Epicypher, version 3), following the user manual version 3.1. Briefly, cells were bound to canavalin-A magnetic beads, permeabilized with 0.01% digitonin, and 5 x 10^5 HC11 cells per target (n=2 per target) were incubated at 4C on a nutator with 0.5 ug of antibody overnight in 50 ul of antibody buffer. Cells were washed and pAG-Mnase was added to the cells in 50 ul of permeabilization buffer and incubated for 10 min at room temperature. Cells were washed again and CaCl2 was added to activate Mnase and incubated at 4C on a nutator for 2 hours. The reaction was stopped with EGTA STOP buffer. *E. coli* DNA was included in the STOP buffer as a spike-in control. Following purification, DNA was quantified using the Qubit dsDNA HS kit. CUT&RUN sequencing libraries were prepared using the CUTANA CUT&RUN Library Prep kit (Epicypher, version 1), following the user manual version 1.4. Briefly, 5 ng of CUT&RUN DNA per target replicate were added at equal volumes for end repair for 20 minutes at 20C. Following adapter ligation for 15 min at 20C and U-excision for 15 min at 37C, DNA was purified using SPRIselect magnetic beads (Beckman Coulter). Next, library amplification was performed for 14 cycles of indexing PCR, followed by PCR cleanup using 1X volume of SPRIselect beads. Sequencing libraries were analyzed by TapeStation (Agilent) and sequenced on the NovaSeq 6000 (Illumina) at 2x50 bp for 8-10 million reads per library.

### CUT&RUN Analysis

Raw FASTQ files were processed using the nf-core CUT&RUN pipeline (nf-core/cutandrun v3.2.2). Adapter and quality trimming was performed using Trim Galore^63^. Post-trim quality control was performed using FastQC^64^. Trimmed FASTQ reads and E. coli spike-in reads were aligned to mm39 and K12-MG1655, respectively, using Bowtie2^65^. Peaks were called using SEACR^66^ and consensus peaks were merged using BEDTools^67^. Heatmaps were made using deepTools^68^. Genomic tracks were visualized using IGV^69,70^.

### RNA Sequencing

Lineage-negative mammary epithelial cells obtained after MACS enrichment were pelleted and resuspended in DNA/RNA Shield reagent (Zymo Research, #R1100) to preserve RNA integrity. Total RNA was purified using the RNeasy Mini Kit (Qiagen, #74104) according to the manufacturer’s protocol. mRNA libraries were generated using the KAPA Stranded mRNA-Seq Kit (Roche/KAPA Biosystems) and assessed for quality and fragment size using an Agilent TapeStation system. Libraries were sequenced on an Element AVITI platform using 2 × 75 bp paired-end chemistry, yielding 27–41 million reads per sample. RNA-seq data processing and analysis are described below.

### RNA Sequencing Analysis

For analysis of public datasets, raw sequencing reads and metadata were obtained from the Gene Expression Omnibus (GEO) using the nf-core/fetchngs pipeline^71^. For all RNA-seq analysis, FASTQ files were processed using the nf-core/rnaseq pipeline^72^. Adapter and quality trimming was performed using Trim Galore^63^. Trimmed FASTQ files were aligned to the mouse GRCm39 reference genome using STAR^73^. RNA sequencing transcripts were quantified using Salmon^74^. Quantified transcripts were analyzed using the nf-core/differentialabundance pipeline^75^. Differential expression analysis was performed using DESeq2^76^ and EnhancedVolcano^77^ from Bioconductor in R^78,79^ RNA sequencing normalized expression plots were made using ggplot2^80^.

### Whole Mount Immunofluorescence

Whole-mount immunofluorescence was performed as previously described (Arora R et al., 2016). GD5.0 E2F5flox/flox and MMTV-cre;E2F5flox/flox mammary glands were dissected from mice and fixed in DMSO:methanol (1:4). Subsequently, they were rehydrated in a methanol:PBST (1% Triton X-100 in PBS) (1:1) solution for 15 minutes, followed by a PBST wash for 15 minutes. Mammary glands were then incubated in a blocking solution (2% powdered milk in PBST) for 1 hour at room temperature. Mammary glands were washed once for 15 minutes with 1% PBST followed by four additional PBST washes for 45 minutes each at room temperature. Mammary glands were then incubated with Hoechst (Sigma Aldrich, B2261) at 4°C for three nights. Samples were washed once for 15 min and three more times for 45 minutes each with 1% PBST, and then stepwise dehydrated into 100% methanol. Mammary glands were incubated overnight at 4°C in a 3% H2O2 solution prepared in methanol. The next day, the samples were washed twice for 15 minutes each and a final 1-hour wash in 100% methanol at room temperature, and then cleared in BABB (1:2, benzyl alcohol:benzyl benzoate) (Sigma-Aldrich, 108006, B6630) overnight.

### Confocal Imaging and Analysis

Whole tissue immunofluorescence samples were imaged using a Leica TCS SP8 X Confocal Laser Scanning Microscope System with white-light laser, 10X air objective and 20X BABB objective. Using the tile scan function with Z stacks 7μm apart (10X), images covering the entire length and thickness of the mammary glands were acquired. Higher resolution images of mammary glands were acquired using a 20X BABB objective. A 4x4 region of mammary gland was captured using the tile scan function, and Z stacks were acquired at 5μm intervals.

Image analysis was performed using commercial software Imaris v9.2.1 (Bitplane). The confocal image (.LIF) files were imported into the Surpass mode of Imaris. 3D images were visualized using the Volume function. Epithelial signal was cleaned by subtracting the CD31+ vessels from the Hoechst or the epithelium-stained channel. Images and movies were captured using the Snapshot and Animation function in Imaris respectively.

### Statistics

For alveoli quantification using pixel density, we performed a two-tailed T-Test. For differential expression analysis, DESeq2 calculates *p*-values using the Wald test and BH-adjusted *p*-value (padj) is calculated using the Benjamini Hochberg method.

### Data Availability

Raw RNA-seq and CUT&RUN data generated for this study are available in the BioProject database under [PRJNA######]. Public datasets were obtained from the Gene Expression Omnibus (GEO) database under the following accession codes: <u>GSE60450</u>, <u>GSE115369</u>, <u>GSE127140</u>, <u>GSE231440</u>.

## Acknowledgements

The authors thank Amy Porter and the Laboratory for Investigative Histopathology at Michigan State University for providing histology services. The authors thank the Van Andel Genomics Core (RRID:SCR_022913) for providing sequencing facilities and services. Illustrations were created using Biorender.com.

## References

1. Macias, H. & Hinck, L. Mammary gland development. WIREs Developmental Biology 1, 533–557 (2012).

2. Robinson, G. W., Karpf, A. B. C. & Kratochwil, K. Regulation of Mammary Gland Development by Tissue Interaction. J Mammary Gland Biol Neoplasia 4, 9–19 (1999).

3. Paine, I. S. & Lewis, M. T. The Terminal End Bud: the Little Engine that Could. J Mammary Gland Biol Neoplasia 22, 93–108 (2017).

4. Shehata, M. et al. Phenotypic and functional characterisation of the luminal cell hierarchy of the mammary gland. Breast Cancer Research 14, R134 (2012).

5. Asselin-Labat, M.-L. et al. Gata-3 Negatively Regulates the Tumor-Initiating Capacity of Mammary Luminal Progenitor Cells and Targets the Putative Tumor Suppressor Caspase-14. Molecular and Cellular Biology 31, 4609–4622 (2011).

6. Watson, C. J. Alveolar cells in the mammary gland: lineage commitment and cell death. Biochem J 479, 995–1006 (2022).

7. Oakes, S. R., Hilton, H. N. & Ormandy, C. J. Key stages in mammary gland development - The alveolar switch: coordinating the proliferative cues and cell fate decisions that drive the formation of lobuloalveoli from ductal epithelium. Breast Cancer Research 8, 207 (2006).

8. Holliday, H., Baker, L. A., Junankar, S. R., Clark, S. J. & Swarbrick, A. Epigenomics of mammary gland development. Breast Cancer Research 20, 100 (2018).

9. Pellacani, D. et al. Analysis of Normal Human Mammary Epigenomes Reveals Cell-Specific Active Enhancer States and Associated Transcription Factor Networks. Cell Reports 17, 2060–2074 (2016).

10. Milevskiy, M. J. G. et al. Three-dimensional genome architecture coordinates key regulators of lineage specification in mammary epithelial cells. Cell Genomics 3, (2023).

11. dos Santos, C. O., Dolzhenko, E., Hodges, E., Smith, A. D. & Hannon, G. J. An Epigenetic Memory of Pregnancy in the Mouse Mammary Gland. Cell Reports 11, 1102–1109 (2015).

12. Bouras, T. et al. Notch Signaling Regulates Mammary Stem Cell Function and Luminal Cell-Fate Commitment. Cell Stem Cell 3, 429–441 (2008).

13. Chakrabarti, R. et al. Elf5 Regulates Mammary Gland Stem/Progenitor Cell Fate by Influencing Notch Signaling. Stem Cells 30, 1496–1508 (2012).

14. Kouros-Mehr, H., Slorach, E. M., Sternlicht, M. D. & Werb, Z. GATA-3 Maintains the Differentiation of the Luminal Cell Fate in the Mammary Gland. Cell 127, 1041–1055 (2006).

15. Asselin-Labat, M.-L. et al. Gata-3 is an essential regulator of mammary-gland morphogenesis and luminal-cell differentiation. Nat Cell Biol 9, 201–209 (2007).

16. Bernardo, G. M. et al. FOXA1 is an essential determinant of ERα expression and mammary ductal morphogenesis. Development 137, 2045–2054 (2010).

17. Fu, N. Y. et al. Foxp1 Is Indispensable for Ductal Morphogenesis and Controls the Exit of Mammary Stem Cells from Quiescence. Developmental Cell 47, 629–644.e8 (2018).

18. Lydon, J. P. et al. Mice lacking progesterone receptor exhibit pleiotropic reproductive abnormalities. Genes Dev. 9, 2266–2278 (1995).

19. Humphreys, R. C., Lydon, J. P., O’Malley, B. W. & Rosen, J. M. Use of PRKO Mice to Study the Role of Progesterone in Mammary Gland Development. J Mammary Gland Biol Neoplasia 2, 343–354 (1997).

20. Mulac-Jericevic, B., Lydon, J. P., DeMayo, F. J. & Conneely, O. M. Defective mammary gland morphogenesis in mice lacking the progesterone receptor B isoform. Proceedings of the National Academy of Sciences 100, 9744–9749 (2003).

21. Fata, J. E. et al. The Osteoclast Differentiation Factor Osteoprotegerin-Ligand Is Essential for Mammary Gland Development. Cell 103, 41–50 (2000).

22. Lee, H. J. et al. Progesterone drives mammary secretory differentiation via RankL-mediated induction of Elf5 in luminal progenitor cells. Development 140, 1397–1401 (2013).

23. Gouilleux, F., Wakao, H., Mundt, M. & Groner, B. Prolactin induces phosphorylation of Tyr694 of Stat5 (MGF), a prerequisite for DNA binding and induction of transcription. The EMBO Journal 13, 4361–4369 (1994).

24. Ormandy, C. J. et al. Null mutation of the prolactin receptor gene produces multiple reproductive defects in the mouse. Genes Dev. 11, 167–178 (1997).

25. Liu, X. et al. Stat5a is mandatory for adult mammary gland development and lactogenesis. Genes Dev. 11, 179–186 (1997).

26. Khaled, W. T. et al. The IL-4/IL-13/Stat6 signalling pathway promotes luminal mammary epithelial cell development. Development 134, 2739–2750 (2007).

27. Liu, X., Gallego, M. I., Smith, G. H., Robinson, G. W. & Hennighausen, L. Functional Rescue of Stat5a-nuIl Mammary Tissue through the Activation of Compensating Signals Including Stat5b. (1998).

28. Oakes, S. R. et al. The Ets transcription factor Elf5 specifies mammary alveolar cell fate. Genes Dev. 22, 581–586 (2008).

29. Nightingale, R. et al. Ehf controls mammary alveolar lineage differentiation and is a putative suppressor of breast tumorigenesis. Developmental Cell 59, 1988–2004.e11 (2024).

30. Nevins, J. R. E2F: a Link Between the Rb Tumor Suppressor Protein and Viral Oncoproteins. Science 258, 424–429 (1992).

31. Dyson, N. The regulation of E2F by pRB-family proteins. Genes Dev. 12, 2245–2262 (1998).

32. Ikeda, M. A., Jakoi, L. & Nevins, J. R. A unique role for the Rb protein in controlling E2F accumulation during cell growth and differentiation. Proceedings of the National Academy of Sciences 93, 3215–3220 (1996).

33. Rempel, R. E. et al. Loss of E2F4 Activity Leads to Abnormal Development of Multiple Cellular Lineages. Molecular Cell 6, 293–306 (2000).

34. Hiebert, S. W. et al. E2F-1:DP-1 Induces p53 and Overrides Survival Factors To Trigger Apoptosis. Molecular and Cellular Biology 15, 6864–6874 (1995).

35. Field, S. J. et al. E2F-1 Functions in Mice to Promote Apoptosis and Suppress Proliferation. Cell 85, 549–561 (1996).

36. Blattner, C., Sparks, A. & Lane, D. Transcription Factor E2F-1 Is Upregulated in Response to DNA Damage in a Manner Analogous to That of p53. Molecular and Cellular Biology 19, 3704–3713 (1999).

37. Höfferer, M., Wirbelauer, C., Humar, B. & Krek, W. Increased levels of E2F-1-dependent DNA binding activity after UV- or γ-irradiation. Nucleic Acids Res 27, 491–495 (1999).

38. Goto, Y., Hayashi, R., Kang, D. & Yoshida, K. Acute loss of transcription factor E2F1 induces mitochondrial biogenesis in Hela cells. Journal of Cellular Physiology 209, 923–934 (2006).

39. Blanchet, E. et al. E2F transcription factor-1 regulates oxidative metabolism. Nat Cell Biol 13, 1146–1152 (2011).

40. Hsu, J. et al. E2F4 regulates transcriptional activation in mouse embryonic stem cells independently of the RB family. Nat Commun 10, 2939 (2019).

41. Andrechek, E. R., Mori, S., Rempel, R. E., Chang, J. T. & Nevins, J. R. Patterns of cell signaling pathway activation that characterize mammary development. Development 135, 2403–2413 (2008).

42. Lindeman, G. J. et al. A specific, nonproliferative role for E2F-5 in choroid plexus function revealed by gene targeting. Genes Dev. 12, 1092–1098 (1998).

43. Gaubatz, S. et al. E2F4 and E2F5 Play an Essential Role in Pocket Protein–Mediated G1 Control. Molecular Cell 6, 729–735 (2000).

44. Ma, L., Quigley, I., Omran, H. & Kintner, C. Multicilin drives centriole biogenesis via E2f proteins. Genes Dev. 28, 1461–1471 (2014).

45. Dagnino, L. et al. Expression patterns of the E2F family of transcription factors during mouse nervous system development. Mechanisms of Development 66, 13–25 (1997).

46. Expression patterns of the E2F family of transcription factors during murine epithelial development. Cell Growth Differ 8, 553–563 (1997).

47. Danielian, P. S., Hess, R. A. & Lees, J. A. E2f4 and E2f5 are essential for the development of the male reproductive system. Cell Cycle 15, 250–60 (2016).

48. To, B. et al. Insight into mammary gland development and tumor progression in an E2F5 conditional knockout mouse model. Oncogene 43, 3402–3415 (2024).

49. Feigman, M. J. et al. Pregnancy reprograms the epigenome of mammary epithelial cells and blocks the development of premalignant lesions. Nat Commun 11, 2649 (2020).

50. Kang, K., Yamaji, D., Yoo, K. H., Robinson, G. W. & Hennighausen, L. Mammary-Specific Gene Activation Is Defined by Progressive Recruitment of STAT5 during Pregnancy and the Establishment of H3K4me3 Marks. Molecular and Cellular Biology 34, 464–473 (2014).

51. Trimarchi, J. M. & Lees, J. A. Sibling rivalry in the E2F family. Nat Rev Mol Cell Biol 3, 11–20 (2002).

52. Yoon, J. et al. E2F and STAT3 provide transcriptional synergy for histone variant H2AZ activation to sustain glioblastoma chromatin accessibility and tumorigenicity. Cell Death Differ 29, 1379–1394 (2022).

53. Zappia, M. P. et al. E2F regulation of the Phosphoglycerate kinase gene is functionally important in Drosophila development. Proceedings of the National Academy of Sciences 120, e2220770120 (2023).

54. Lodewijk, G. A. et al. E2F1 induces a G0-G1 reentry transcriptional program without changing chromatin accessibility. 2025.09.23.678145 Preprint at 10.1101/2025.09.23.678145 (2025).

55. Lim, E. et al. Transcriptome analyses of mouse and human mammary cell subpopulations reveal multiple conserved genes and pathways. Breast Cancer Res 12, R21 (2010).

56. Bracken, A. P. et al. EZH2 is downstream of the pRB-E2F pathway, essential for proliferation and amplified in cancer. The EMBO Journal 22, 5323–5335 (2003).

57. Yang, R. et al. E2F7−EZH2 axis regulates PTEN/AKT/mTOR signalling and glioblastoma progression. Br J Cancer 123, 1445–1455 (2020).

58. Regan, J. L. et al. c-Kit is required for growth and survival of the cells of origin of Brca1-mutation-associated breast cancer. Oncogene 31, 869–883 (2012).

59. Dawson, C. A. et al. Hormone-responsive progenitors have a unique identity and exhibit high motility during mammary morphogenesis. Cell Reports 43, 115073 (2024).

60. Chen, C.-R., Kang, Y., Siegel, P. M. & Massagué, J. E2F4/5 and p107 as Smad Cofactors Linking the TGFβ Receptor to c-*myc* Repression. Cell 110, 19–32 (2002).

61. Blakely, C. M. et al. Developmental stage determines the effects of MYC in the mammary epithelium. Development 132, 1147–1160 (2005).

62. To, B. & Andrechek, E. R. Transcription factor compensation during mammary gland development in E2F knockout mice. PLOS ONE 13, e0194937 (2018).

63. Krueger, F., et al. FelixKrueger/TrimGalore: v0.6.10 - add default decompression path. Zenodo 10.5281/zenodo.5127898 (2023).

64. Andrews, S. FastQC: a quality control tool for high throughput sequence data.

65. Langmead, B. & Salzberg, S. L. Fast gapped-read alignment with Bowtie 2. Nat Methods 9, 357–359 (2012).

66. Meers, M. P., Tenenbaum, D. & Henikoff, S. Peak calling by Sparse Enrichment Analysis for CUT&RUN chromatin profiling. Epigenetics & Chromatin 12, 42 (2019).

67. Quinlan, A. R. & Hall, I. M. BEDTools: a flexible suite of utilities for comparing genomic features. Bioinformatics 26, 841–842 (2010).

68. Ramírez, F. et al. deepTools2: a next generation web server for deep-sequencing data analysis. Nucleic Acids Res 44, W160–W165 (2016).

69. Robinson, J. T. et al. Integrative genomics viewer. Nat Biotechnol 29, 24–26 (2011).

70. Thorvaldsdóttir, H., Robinson, J. T. & Mesirov, J. P. Integrative Genomics Viewer (IGV): high-performance genomics data visualization and exploration. Brief Bioinform 14, 178–192 (2013).

71. Patel, H., et al. nf-core/fetchngs: nf-core/fetchngs v1.12.0 - Titanium Platypus. Zenodo 10.5281/zenodo.5070524 (2024).

72. Patel, H., et al. nf-core/rnaseq: nf-core/rnaseq v3.22.0 - Palladium Penguin. Zenodo 10.5281/zenodo.1400710 (2025).

73. Dobin, A. et al. STAR: ultrafast universal RNA-seq aligner. Bioinformatics 29, 15–21 (2013).

74. Patro, R., Duggal, G., Love, M. I., Irizarry, R. A. & Kingsford, C. Salmon provides fast and bias-aware quantification of transcript expression. Nat Methods 14, 417–419 (2017).

75. Wacker O et al. nf-core/differentialabundance: v1.4.0 - 2023-11-27. Zenodo 10.5281/zenodo.7568000 (2023).

76. Love, M. I., Huber, W. & Anders, S. Moderated estimation of fold change and dispersion for RNA-seq data with DESeq2. Genome Biol 15, 550 (2014).

77. Blighe, K. kevinblighe/EnhancedVolcano. (2025).

78. Gentleman, R. C. et al. Bioconductor: open software development for computational biology and bioinformatics. Genome Biol 5, R80 (2004).

79. Team, R. C. R: A language and environment for statistical computing. R Foundation for Statistical Computing, Vienna, Austria (2021).

80. ggplot2 - Wickham - 2011 - WIREs Computational Statistics - Wiley Online Library. https://wires.onlinelibrary.wiley.com/doi/full/10.1002/wics.147.

